# Probing dynamical cortical gating of attention with concurrent TMS-EEG

**DOI:** 10.1101/615708

**Authors:** Yuka O. Okazaki, Yuji Mizuno, Keiich Kitajo

**Affiliations:** CBS-TOYOTA Collaboration Center, RIKEN Center for Brain Science, 2-1 Hirosawa, Wako, Saitama, 351-0198, Japan; Research Fellow of Japan Society for the Promotion of Science (JSPS), 5-3-1 Kojimachi, Chiyoda-ku, 102-0083, Tokyo, Japan; Division of Neural Dynamics, Department of System Neuroscience, National Institute for Physiological Sciences, National Institutes of Natural Sciences, 38 Nishigonaka, Myodaiji, Okazaki, Aichi, 444-8585, Japan; Department of Physiological Sciences, School of Life Science, The Graduate University for Advanced Studies (SOKENDAI), 38 Nishigonaka, Myodaiji, Okazaki, Aichi, 444-8585, Japan

**Keywords:** TMS-EEG, visual attention, cortical excitability, effective connectivity, alpha oscillations

## Abstract

**Background:** Attention facilitates gating of information from the sending brain area to receiving areas, with this being achieved by dynamical change in effective connectivity between cortices. However, it is difficult to assess effective connectivity, which refers to the causal influence of one cortical area on another.

**Objective:** The aim of this study is to directly probe the effective connectivity between cortical areas, which is modulated by covertly shifted attention, excluding the thalamic influence.

**Methods:** Transcranial magnetic stimulation (TMS) was used to directly perturb the right retinotopic visual cortex in task-relevant (TR) and task-irrelevant (TIR) networks, and the impact of this was tracked to other areas by concurrent use of electroencephalography (EEG).

**Results:** Key finding was that TMS to the TR hemisphere led to a stronger evoked potential than did
stimulation to the TIR hemisphere. Moreover, stronger beta- and gamma-band effective connectivities assessed as lagged phase synchronizations between stimulated areas and other areas were observed when TMS was delivered to the TR area. These effects were more enhanced when they preceded more prominent alpha lateralization, which is known to be associated with gating information.

**Conclusion:** Our results indicate that attention-regulated cortical excitability and feedforward dynamical effective connectivity can be probed by direct cortical stimulation to the TR or TIR area, thereby bypassing thalamic gating. These results bear out the idea that TMS-EEG could help to characterize changes in the functional architecture of brain networks that are required for rapid adaptation to the environment or due to brain diseases.

**Highlights:**

- Attention-regulated cortical excitability and effective connectivity were probed by TMS-EEG.
- Stronger TMS evoked potentials were observed in the task-relevant visual hemisphere.
- Strong effective connectivity between task-relevant visual cortex and other areas were found.

## Introduction

In our daily lives, we are bombarded with sensory inputs far beyond our information processing capacity. The covert direction of attention to specific parts of a visual scene allows processing resources to be allocated to potentially relevant stimuli, typically through the modulation of local neuronal excitability to attended and unattended sensory inputs [1–4].

However, attention can also change signal transmission, which is mediated by large-scale phase synchronizations in neural activity between task-relevant regions [5–9]. In a study using a covert visual attention paradigm, Doesburg and colleagues showed that gamma-band synchronization between the contralateral occipital electrode and other electrodes increased during attention maintenance [5, 10]. Their results suggested that long-range gamma synchronization helps establish a transient network that promotes information transmission from modality-specific cortical areas to other cortical areas forming TR networks. On one hand, synchronizations between scalp electrodes can be a result of spurious coupling that is actually driven by a common source or contamination from volume conduction [11, 12], while on the other hand, the causal influence of one brain region on others has been assessed by measuring effective connectivity [13, 14]. The most straightforward way to measure effective connectivity is to perturb a part of the brain network and observe how its impact is transmitted to other sites. In human studies, this has been achieved by combining transcranial magnetic stimulation (TMS) with EEG [13–19]. TMS-EEG has allowed demonstration of the dynamical properties of effective connectivity by showing how the propagation patterns of TMS evoked potentials could be used to differentiate between sleep and wakeful states [13], as well as the propagation of TMS-induced transient phase resetting of ongoing oscillations from visual to motor areas [17]. In this study, we used such a perturbation approach to probe the dynamical nature of cortical excitability and effective connectivity alterations between different attention conditions. The spatiotemporal profiles of EEG responses were examined when TMS was applied to the right V1/V2 of the task-relevant (TR) or task-irrelevant (TIR) hemisphere, depending on the attention direction.

EEG-level phase synchronization between distant areas may be a plausible mechanism for network communication [20, 21]. If attention modulates effective connectivity depending on the task at hand, we hypothesize that effective connectivity assessed as lagged phase synchronizations would increase between TR areas when one of these areas is perturbed, while perturbation to TIR areas would not induce such a prominent change in phase dynamics because effective connectivity would be decreased. In addition, such modulation of effective connectivity may depend on the extent to which alpha oscillations are modulated in the preparatory period [22].

## Materials and methods

### Participants

Twenty-two healthy right-handed participants (7 female and 15 male, mean age: 24.9 ± 5.7 [SD]) gave informed written consent for their participation in this study. The ethical committee of the RIKEN Center for Brain Science approved this TMS-EEG study.

### Stimulus and task

The TMS-EEG experiment consisted of six runs (three runs for real TMS, and three runs for
sham TMS), each containing 96 trials. Participants were seated 100 cm from a gamma corrected
LCD monitor (BenQ XL2420, 100 Hz refresh rate) and performed a cued spatial attention task during both the real- and sham-TMS runs, which were performed in random order (Fig. 1a). A trial began with an arrow cue (0.1 s) indicating the hemifield to which the participant should attend, followed by an anticipatory interval of 1.2 s. Subsequently, either a bilateral visual stimulus was presented or TMS was applied to the right V1/V2 area (see EEG recordings and TMS), with the order of these being randomly determined. In the visual stimulation trials, a target Gabor grating (standard deviation of the Gaussian envelope, 0.18; spatial frequency, 2.5 cycles per degree [cpd]; contrast, 50%; orientation, ± 2°) was presented in the cued hemifield together with a distractor Gabor grating (orientation, ± 45°) in the other hemifield. These grating stimuli were presented for 0.05 s, followed by a 0.05 s bilateral backward mask stimulus (radial Gabor grating with the same property as the target). Then, after 1 s, participants were required to indicate whether the target stimulus was tilted to the right or left by pressing the arrow keys with the index or middle finger of their dominant right hand for the left or right orientation, respectively, with a response interval of 2 s being allowed for this, during which the color of a fixation cross changed from white to green. In the TMS trials with no visual stimulation, the participants were asked to press freely either the left or right arrow key during the response interval. Because it was unpredictable and counterbalanced as to whether visual stimuli or TMS would be applied, the participants had to attend to the cued direction in both conditions. Therefore, in TMS trials, either the TR or TIR right hemisphere could be perturbed, depending on the attended direction (Fig. 1b). Stimulus delivery was controlled using Psychtoolbox-3 [23–25].

**Figure 1.**
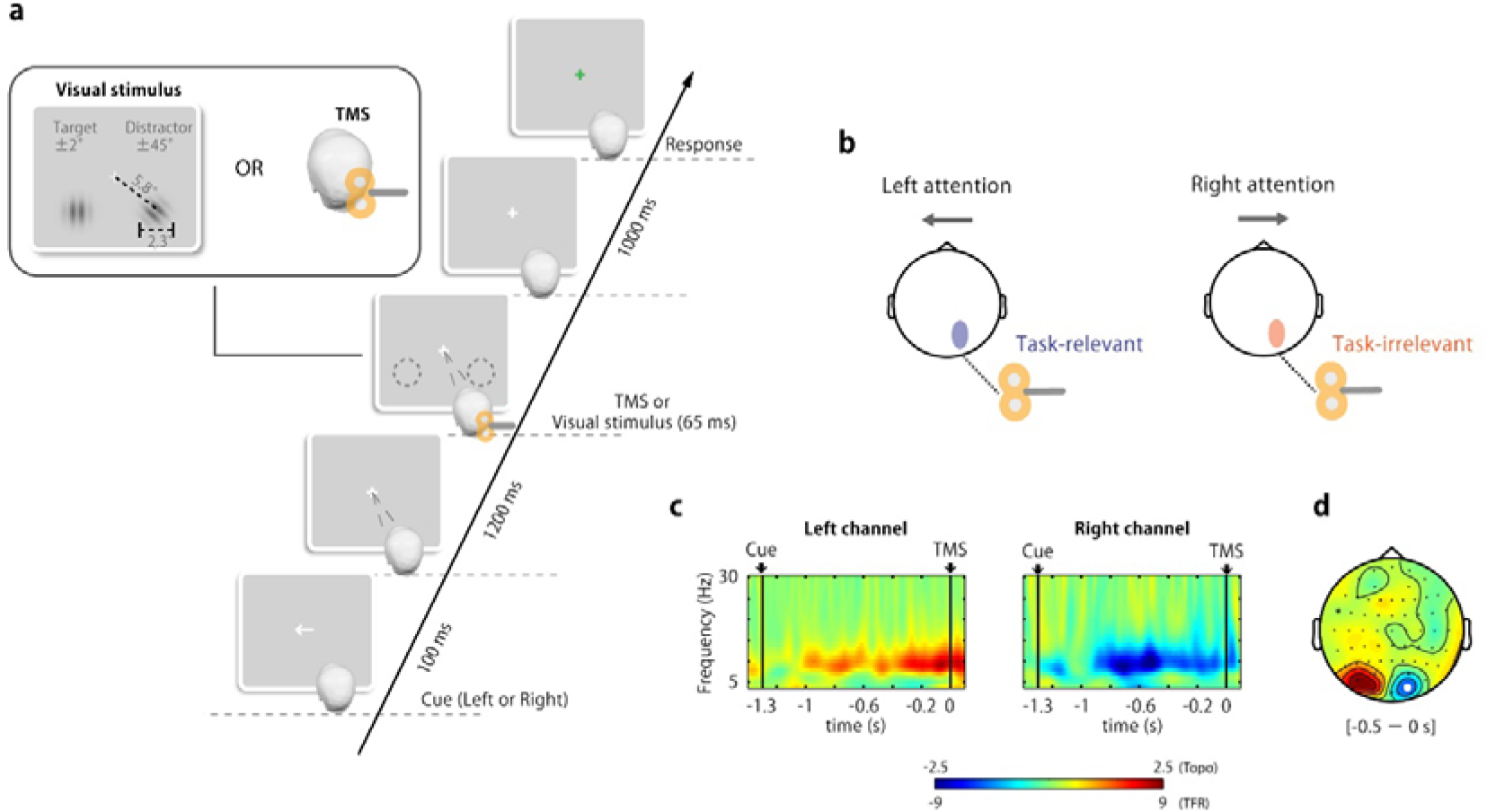
**(a)** Experimental paradigm. The trial started with a cue indicating which hemifield to attend to, and after 1.3 s, TMS was applied to the right visual cortex without presentation of a visual stimulus. To guarantee the participant’s attention to the cued hemifield, trials requiring a response to the target Gabor orientation in the cued hemifield were randomly introduced. The target Gabor stimulus (± 2° oriented) in the cued side was always presented with a distractor stimulus (± 45° oriented) on the opposite side. Participants were unable to predict whether a visual stimulus or TMS would be applied, so they needed to follow the cued instructions. **(b)** A schematic figure for the attention-dependent conditions. The right visual cortex where TMS was applied becomes the task-relevant hemisphere in the left attention trials, while it becomes the task-irrelevant hemisphere in the right attention trials. **(c)** Grand averaged time-frequency representations of the AMI (left cue - right cue) in the left and right electrode. **(d)** Grand averaged topographic AMI (8–12 Hz) for the time range from −0.5 to 0 s. The white dot indicates the electrode position near the TMS coil.

### EEG recordings and TMS

EEG (left earlobe reference; ground AFZ) signals were recorded from 63 scalp sites using sintered Ag/AgCl TMS-compatible electrodes mounted on a 10/10 system EasyCap (EASYCAP Gmbh, Germany). Horizontal and vertical electrooculography (EOG) signals were continuously recorded. The electrode impedance was kept below 10 kΩ. The EEG and EOG signals were amplified and recorded by a Brain Amp MR+ (Brain Products, Germany) system with a sampling rate of 5 kHz. The electrode lead wires were arranged orthogonal to the TMS coil handle direction, to reduce TMS-induced artifacts [26]. The TMS target site was located at the upper right V1/V2 according to the calcarine sulcus determined on each individual subject’s MRI (mean Talairach coordinates ± SD: 11, −95 ± 3, 5), and the TMS coil position was eventually near to the O2 electrode for all participants. This TMS location approximately corresponds to the alpha modulated area (see Fig. 1c). The TMS coil and head position were continuously monitored using Brainsight TMS (Rogue Research Inc., Canada), and kept within 5 mm of the initial position. For the sham stimulation, the coil was rotated 90° around the handle axis and spaced from the head using a 3 cm plastic cube [27]. Thus, the participant received some sensation of vibration caused by the TMS click, without receiving direct cortical stimulation. Additionally, the click sound was attenuated by the participant wearing earplugs and the delivery of white masking noise in all conditions. The stimulation intensity achieving a 95% active motor threshold in the right first dorsal interosseous muscle was individually adjusted according to the distance between the TMS coil and targeted visual cortex [28].

### TMS and ocular artifact rejections

EEG data were analyzed using in-house developed scripts written in MATLAB (MathWorks, Natick, USA) and FieldTrip [29]. The data were first segmented into 5 s epochs (3.5 s pre-stimulation and 1.5 s post-stimulation), and then the epoched data were re-referenced offline to the average of the right and left earlobe signals. TMS and ocular artifacts were rejected using the following steps. First, the data samples were temporally smoothed using linearly interpolation (two samples preceding and about 21 samples following the TMS onset, i.e., 4.3 ± 0.08 ms) to remove excessive TMS artifacts. Second, epochs contaminated with eye movements and blinks in the interval between −1.3 and 1.1 s were discarded according to the following criteria: horizontal EOG signals exceeding ~50 μV, which approximately corresponded to the stimulus eccentricity (6° visual angle), and vertical EOG exceeding ~100 μV. Third, the exponential decay artifacts due to TMS were attenuated using independent component analysis (ICA) based on the method proposed by Korhonen and colleagues [30] (see also http://www.fieldtriptoolbox.org/tutorial/tms-eeg for a more practical application). ICs were excluded according to mean z-score values greater than 1.65 between 0 and 50 ms, with the topography of the mixing matrix being confined within the stimulated region. For ocular artifacts, ICs correlating with the EOG (*r* > 0.2, *p* < 0.05) and showing the typical topographical structure of saccadic eye movements and blinks were also excluded. Finally, temporal smoothing linear interpolation was again applied until 10 ms to remove residual artifact. Four participants for whom more than 45% of trials were lost after discarding epochs contaminated with ocular artifacts were excluded from further analysis. The artifact-removed EEG measures were then down-sampled to 500 Hz. Current source density (CSD) transformation [31, 32] was applied to localize the activity and attenuate the effects of volume conduction.

### TMS evoked potential (TEP)

To investigate the local cortical excitability, we took particular note of an initial cortical reaction in TEP. The preprocessed EEG epochs were bandpass filtered from 3 to 45 Hz using a fourth order Butterworth filter and averaged for each condition. The peak TEP components in the interval 0-0.2 s were identified.

### Attentional alpha power modulation

The instantaneous amplitude and phase were obtained using a wavelet transform with a center frequency *f*, time *t*, and standard deviation σ_*f*_ = *4f/m* and σ_*t*_ = *m/2πf* [33]. The constant *m* was set to 3. Next the alpha modulation index was computed according to the following: *AMI*_*ch*_ = *α*_*ch (left cue)*_ − *α*_*ch (right cue)*_, where *α*_*ch (left cue)*_ and *α*_*ch (right cue)*_ refer to the mean alpha power computed from the instantaneous amplitude for the left and right cued trials per electrode. For each participant, the most positive electrode in the left hemisphere and the most negative electrode in the right hemisphere at the time when the AMI was topographically maximized between −1 and −0.1 s was chosen. The grand averaged AMI and TFR of power from an individually selected electrode and topography are shown in Figure 1c and d, respectively. Using selected electrodes, the alpha lateralization index was computed as *ALI* = *α*_*ipsilateral electrode*_ − *α*_*contralateral electrode*_, which represents the contrast in alpha power between the ipsilateral and contralateral electrode in respect to the attention side. Then, the 50% of trials with the highest ALI were assigned to “high-ALI” trials, while the other half of trials were assigned to “low-ALI” trials. This meant that the alpha power in the TMS target area (right V1/V2) was relatively low in the left cue trials and high in the right cue trials in the high-ALI trials, while this contrast was marginal in the low-ALI trials.

### Inter-areal phase synchronization

The effective connectivities between O2 and all other electrodes were assessed using the lagged phase locking value to determine the relationship between sending and receiving areas:

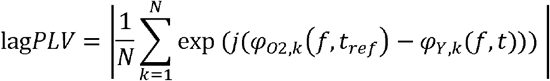

where *φ*_O2_ and *φ*_Y_ are the instantaneous phase of frequency *f* at electrode O2 and electrode *Y* (all the other electrodes). *N* denotes the number of trials, and the reference time was set to *t*_*ref*_ = 0, which was the onset timing of the TMS. Thus, lagPLV evaluates the consistency of the phase difference between electrode O2 (sending) at the TMS onset and the other electrodes (receiving) with a given lag time.

### Statistics

The TEP and lagPLV observed in the TR and TIR conditions were statistically evaluated. Note that these were obtained by subtracting sham conditions to reduce the osteoconductive auditory component.

For the initial cortical reaction to the TMS perturbation, the P20 component (average of five samples) of the O2 electrode was statistically assessed using a two-way ANOVA with task relevancy (TR and TIR) and ALI (low ALI and high ALI) as fixed factors.

For lagPLV, a cluster-based permutation test [34] was used to verify whether visual spatial attention changed the degree of signal transmission from early visual cortex to other cortical areas. A cluster-based permutation test was used to determine whether the observed difference in the cluster statistics between conditions was large enough to reject the null hypothesis, according to the following procedures. First, all elements (i.e., 63 electrodes, 3-45 Hz, 0-0.5 s) in the lagPLV matrices between the left and right attention conditions in each low-ALI and high-ALI set were compared using two-tailed paired *t*-tests. Then, contiguous negative and positive clusters in the matrices were identified according to an uncorrected *p*-value threshold of < 0.05, and the sums of the *t*-values in clusters were calculated as cluster statistics. Second, to obtain the null distribution of the test cluster statistic, the maximal cluster from matrices of randomly permuted two-condition labels within participants was identified, with 500 iterations being used. Finally, using the 97.5 percentile of the null distribution as the level of statistical significance, significant clusters of observed data were identified.

## Results

### Pre-stimulus alpha power modulation by attention

The TFR of the power from individually selected electrodes was contrasted between left and right cue trials and averaged over participants. The grand averaged TFR of the power indicated that alpha power was continuously modulated in the interval between cue and TMS, approximately 1 s prior to the stimulus onset, and corresponding to the direction of attention (Fig. 1c). This result is evidence that the participants were engaged in the attentional task directing their attention toward the visual hemifield of the cue side, even in those TMS trials without visual stimuli, although the behavioral performance was relatively low (correct rate mean±sd: 0.65±0.11). Topographic representation of the mean AMI from −0.5 to 0 s indicates that the alpha power was clearly lateralized in the occipito-parietal electrodes, i.e., the alpha power in the ipsilateral hemisphere (TIR hemisphere) to the attention direction increased, and alpha power in the contralateral hemisphere (TIR hemisphere) decreased (Fig. 1d). Note that the TMS target (around O2) coincides with the region where alpha power was strongly modulated.

### Alpha lateralization-dependent early TMS evoked potential

The TEP was calculated to examine the response to the TMS applied to the target area, with the alpha power just before stimulation being strongly modulated by attention. We identified several TEP components from the O2 electrodes (near to the stimulation site), including the P20, N50, P70, N100, and P120 components, in both the high-ALI and low-ALI trials (Fig. 2a). Cortical excitability influenced by the alpha oscillations should be reflected in an immediate response to the TMS perturbation, e.g., the P20. In high-ALI trials (left panel), the topographical map contrasting the left and right cue trials indicated that P20 showed a maximal difference at approximately the stimulation site, while the maximum differences in low-ALI trials were sparsely distributed. We quantified the effect of pre-stimulus alpha power on the P20 using a two-way ANOVA with the factors ALI type and task relevancy (Fig. 2b). This showed a main effect for task relevance (*F*(1, 17) = 8.756, *p* = 0.009), indicating that the P20 in the TR condition was larger than in the TIR condition, regardless of the amount of alpha lateralization. There was no significant main effect for ALI type (*F*(1, 17) = 1.099, *p* = 0.309) or the interaction between factors (*F*(1, 17) = 0.587, *p* = 0.454). However, we further applied an ad hoc simple main-effect test to examine the difference between TR and TIR, doing this separately for the high-ALI and low-ALI trials. This revealed that the effect of task relevancy on the P20 component was significant in high-ALI trials (*F*(1, 17) = 5.431, *p* = 0.032), but not in low-ALI trials (*F*(1, 17) = 1.398, *p* = 0.253). These results indicate that the hemisphere with task relevance to the attention was in a highly excitable state, but that this was likely to depend on the degree of alpha power in that hemisphere.

**Figure 2.**
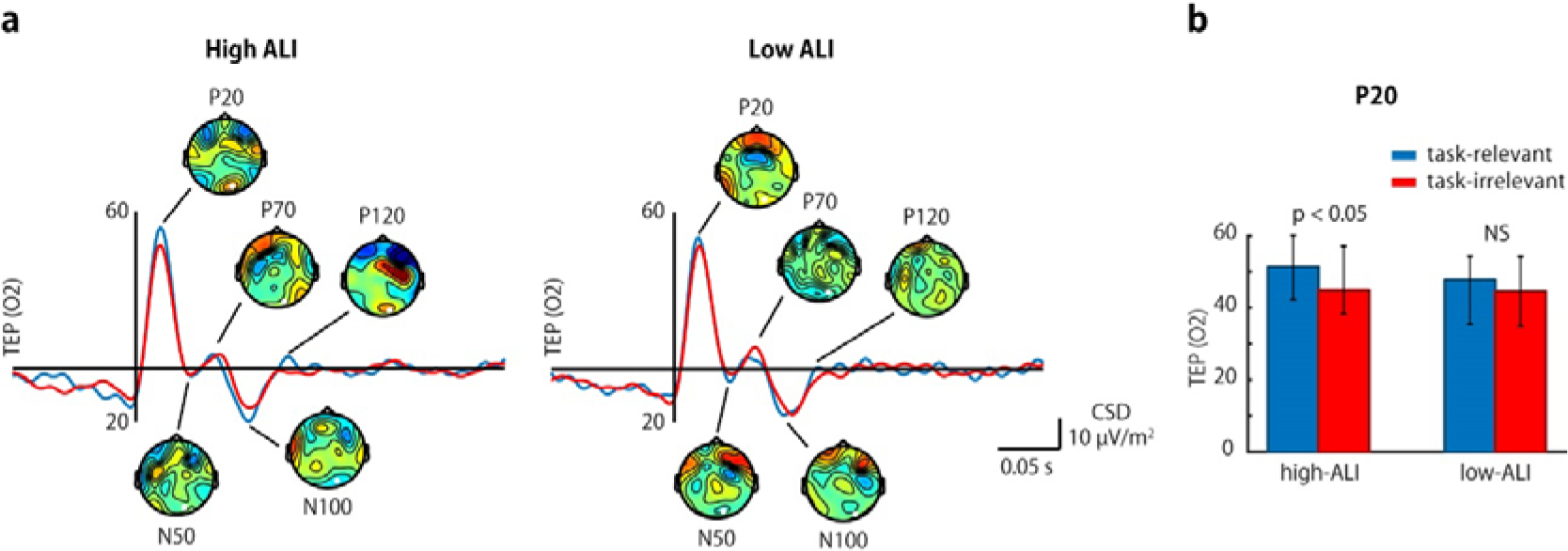
Attentional modulation in TMS evoked potentials. **(a)** TEPs from O2 at the site of stimulation for the high-ALI and low-ALI trials. Topographic differences between the left and right cue trials are shown for each TEP component. **(b)** The first response to TMS, i.e., P20, was significantly larger in the task-relevant condition with high alpha power modulation (high-ALI trials) than in the task-irrelevant condition. The same contrasts with low-ALI trials were negligible.

### Alpha lateralization-dependent effective connectivity

To probe how effective connectivity varies with attention, we assessed lagPLV to estimate the effective connectivity as a directional phase coupling between the sending and receiving areas. LagPLV makes it possible to know when and where the perturbed phase dynamics at one region affect other regions in the brain network. We compared lagPLVs between TR and TIR conditions and observed a significant difference only in the high-ALI trial where the alpha power of the stimulation region was low under the TR condition and high under the TIR condition. There were no significant differences in the low-ALI trials. To quantify the time-frequency characteristics of this significant difference, we examined the number of significant electrode pairs over time and frequency range (Fig. 3a). In the beta and low gamma band with a peak at 25 Hz, the lagPLV of the TR condition was found to be significantly stronger at more electrode pairs than in the TIR condition (*p* < 0.05). The significant difference started at about 70 ms, peaked at 114 ms, and ended at about 150 ms. These results show that the TMS perturbation to the TR region more strongly and widely affected other regions in comparison with the perturbation to the TIR region.

**Figure 3.**
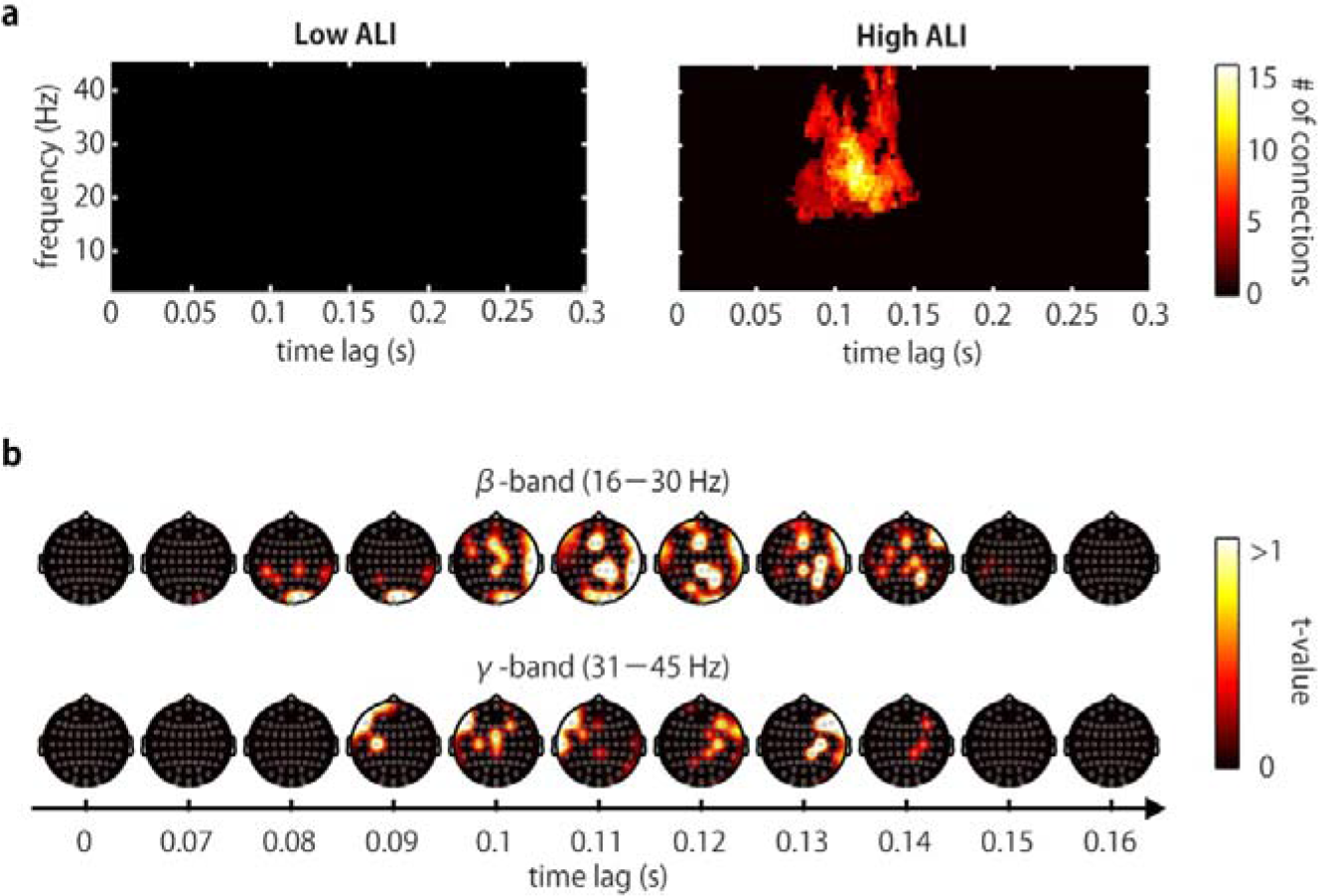
Attentional modulation in effective connectivity from V1/V2 to other areas. **(a)** A time-frequency profile of the number of significant lagPLVs in the comparison between the task-relevant (TR) and task-irrelevant (TIR) conditions indicates that large areas are synchronized in the beta and gamma bands. This significant difference was observed only in high-ALI trials (right panel). **(b)** A spatiotemporal profile of the significant difference between TR and TIR in high-ALI trials shows that TMS perturbation on the task-relevant area has strong impacts on other cortical areas, e.g., parietal and frontal regions.

Next, we investigated the spatial extent of this difference in the beta (16-30 Hz) and gamma bands (31–45 Hz). Figure 3b shows a topographical map of the mean significant *t*-values identified by the cluster-based permutation test. For the difference between the TR and TIR conditions in the beta-band lagPLV, the effect of TMS perturbation on the TR region was confined to the temporal-parietal region until about 100 ms, and then to the frontal region. For the gamma-band lagPLV, the difference was first observed in the frontal region contralateral to the stimulated site, and then moved to the ipsilateral frontal region. These results suggest that networks between short-distance TR areas are established with beta-band oscillations and that the chained response was triggered by a local stimulation, whereas long distance networks may be established with gamma-band oscillations.

## Discussion

### Probing cortical excitability

It is generally agreed that directing attention toward a specific location in the visual field facilitates perception and retinotopic neural responses to subsequently presented target stimuli at the attended location [35, 36]. However, given the evidence from human fMRI studies that attention modulates activity in the lateral geniculate nucleus [37, 38], the enhancement of visual evoked activity in the striate [36, 39] and extrastriate [40, 41] cortices probably reflects amplified afferent inputs through thalamic gating. Thus, it is hard to separate the contribution from the thalamus when evaluating top-down modulation on the neocortex. Bestmann and colleagues tested this issue by direct cortical stimulation of the visual cortex, so that the visual stimulus did not pass through the retinogeniculate pathway [42]. They showed that spatial attention facilitates awareness of phosphenes induced by TMS, although there is still controversy over the relationship between initial sensory responses and behavioral perception.

In an analogous spatial attention paradigm, we observed neural responses to TMS in early visual cortex, instead of phosphene perception. In the current study, the direct neural responses at a very early latency, i.e., P20, were modulated by the direction of attention and its associated alpha power modulation preceding the neural responses. These findings are compatible with a report by Herring and colleagues [43], although in their study, the earliest attentional modulation occurred in the N40 TEP component, rather than in P20. N40 may reflect inhibitory feedback in response to TMS-induced cortical excitation, rather than the initial excitation itself [44]. A plausible explanation for the difference in timing of the attentional effect on the early TEP component may be partly related to the stimulated area. We confined the TMS target area to the right V1/V2, while Herring et al. defined a target area to ensure retinotopic phosphene perception. Given that it has been reported that stimulation sites generating retinotopic phosphenes are situated at V2 and V3 [45], the TEP components observed in Herring’s study might be a direct response from the extrastriate cortex, rather than the striate cortex. Such a hierarchical difference may lead to a difference in timing. In summary, these results provide evidence that the cortico-cortical top-down influence on the excitability of early visual cortices can be regulated in parallel with thalamic gating. Moreover, it seems that a large alpha lateralization emphasizes this attentional effect, i.e., the smaller P20 can be accompanied by large-power alpha oscillations in the TMS target site, and vice versa.

### Probing cortical effective connectivity

The function of attention is not only to change local neural excitability; it must also facilitate dynamical interactions between different areas, depending on the task at hand. This could be achieved by flexible control of effective connectivity mediated by phase synchronization of neural oscillations at the regions sending and receiving information. Doesburg and colleagues showed that TR early visual cortex and other regions were coupled with gamma-band phase synchronization during covertly sustained attention [10]. To demonstrate that such phase synchronization reflects the directional network for signal transmission modulated by attention, we locally perturbed either contralateral or ipsilateral visual cortex during covert attention. In direct perturbation approaches, effective connectivity has been evaluated by measuring the propagation of evoked activity [13, 15]. Although the validity of the lagPLV used here to investigate the effective connectivity is comparable to that of the techniques used in these previously mentioned studies, the lagPLV with TMS perturbation is more focused on directional interactions. When waves originating in the sending area reach the receiving area, the phase difference between these areas becomes constant at that time; that is, it becomes synchronized, but with a certain delay time. We found that the waves in the right early visual area propagated widely and reached to the other cortical areas at certain delay time in the TR condition, while it seemed to be suppressed in the TIR condition. These results suggest that cortical gating of the feedforward input is achieved by regulating the effective connectivity in the phase dynamics between cortical areas. Importantly, it should be noted that successful gating depended on the degree of alpha lateralization caused by attention. Although the mechanism behind the alpha contribution and its source is still unknown, the key role in generating the alpha oscillations may be played by inhibitory GABAergic neurons [46–48]. In a model to test the cortical gating mechanism to TMS evoked activity during slow wave sleep, increased GABA release from local inhibitory neurons in the cortex was effective in reducing the propagation of evoked activity [49]. Thus, the increased effective connectivity in the present study may be mediated by a GABAergic inhibitory feedback in the neocortex or thalamus. We recommend that studies employing magnetic resonance spectroscopy should be performed to reveal the roles of GABA release.

### Effect of the auditory artifact from the TMS click sound

In general, a limitation of TMS research is that the TMS pulse is accompanied by a click sound of about 100 to 120 dB [50], as well as the introduction of several electromagnetic and/or muscle artifacts. The click sound contaminates part of the TEP with an auditory evoked potential (AEP) [51–53]. We used a combination of earplugs and white noise to mask the click sound, and placed a thin layer of foam between the TMS coil and the EEG cap to attenuate bone conduction of sound [13, 53–55]. In addition, we arranged the electrode leads to minimize electromagnetic artifacts during the experiment [26], and attenuated any such artifacts using offline ICA analysis [30, 43]. If phase shifts are caused by an AEP and/or TMS artifacts, we should acknowledge that the lagPLV may spuriously increase, but this would not explain our results. Because the effects of these artifacts should be identical across different attention conditions and we always made comparisons between conditions with sham subtraction, we believe that the modulated lagPLV is associated with dynamical gating of cortical information processing. Nevertheless, it should be noted that further developments in more realistic sham stimulation [56–59] and electrode referencing methods [60] are awaited to allow the TMS-EEG community to maximize the direct effects of TMS on cortical responses.

## Conclusions

By directly stimulating TR or TIR areas, we succeeded in probing attention-regulated cortical excitability and feedforward effective connectivity in the phase dynamics between cortical areas. This method provides evidence that attentional top-down control not only coordinates the sensory input to the cortex by thalamic gating, but also changes the cortical excitability and effective connectivity that mediates cortical gating between TR regions. The observation of perturbation effects caused by direct use of TMS-EEG will allow the characterization of excitability and effective connectivity changes according to the task at hand as well as changes due to diseases which impair networks such as stroke [61].

## Abbreviations

TR: task-relevant
TIR: task-irrelevant
CSD: current source density
TEP: TMS evoked potential
TFR: time-frequency representations
AMI: alpha modulation index
ALI: alpha lateralization index
lagPLV: lagged phase locking value
AEP: auditory evoked potential

## Acknowledgments

This study was supported by JST PRESTO, MEXT Grants-in-Aid for Scientific Research 26282169 and 15H05877, Grant-in-Aid for JSPS Fellows (26-8352), and a research grant from TOYOTA Motor Corporation. We are grateful to Yoko Noguchi for help with TMS-EEG recording.

